# New genetic variants associated with major adverse cardiovascular events in patients with acute coronary syndromes and treated with clopidogrel and aspirin

**DOI:** 10.1101/411165

**Authors:** Xiaomin Liu, Hanshi Xu, Huaiqian Xu, Qingshan Geng, Wai-Ho Mak, Fei Ling, Zheng Su, Fang Yang, Tao Zhang, Jiyan Chen, Huanming Yang, Jian Wang, Xiuqing Zhang, Xun Xu, Huijue Jia, Zhiwei Zhang, Xiao Liu, Shilong Zhong

## Abstract

**Importance:** Although a few studies have reported the effects of several polymorphisms on major adverse cardiovascular events (MACE) in patients with acute coronary syndromes (ACS) and those undergoing percutaneous coronary intervention (PCI), these genotypes account for only a small fraction of the variation and evidence is insufficient. This study aims to identify new genetic variants associated with MACE by large-scale sequencing data.

**Objective:** To identify the genetic variants that caused MACE.

**Design:** All patients in this study were allocated to dual antiplatelet therapy for up to 12 months and have the follow-up duration of 18 months.

**Setting:** A two-stage association study was performed.

**Participants:** We evaluated the associations of genetic variants and MACE in 1961 patients with ACS undergoing PCI (2009-2012), including high-depth whole exome sequencing of 168 patients in the discovery cohort and high-depth targeted sequencing of 1793 patients in the replication cohort.

**Main Outcomes and Measure:** The primary clinical efficacy endpoint was the major adverse cardiovascular events (MACE) composite endpoint, including cardiovascular death, myocardial infarction (MI), stroke (CT or MR scan confirmed) and repeated revascularization (RR).

**Results:** We discovered and confirmed six new genotypes associated with MACE in patients with ACS. Of which, rs17064642 at *MYOM2* increased the risk of MACE (hazard ratio [HR] 2.76; P = 2.95 × 10^-9^) and reached genome-wide significance. The other five suggestive variants were *KRTAP10-4* (rs201441480), *WDR24* (rs11640115), *ECHS1* (rs140410716), *AGAP3* (rs75750968) and *NECAB1* (rs74569896). Notably, the expressions of *MYOM2* and *ECHS1* are down-regulated in both animal models and patients with phenotypes related to MACE. Importantly, we developed the first superior classifier for predicting MACE and achieved high predictive accuracy (0.809).

**Conclusions and Relevance:** We identified six new genotypes associated with MACE and developed a superior classifier for predicting MACE. Our findings shed light on the pathogenesis of cardiovascular outcomes and may help clinician to make decision on the therapeutic intervention for ACS patients.

**Trial Registration:** This study has been registered in the Chinese Clinical Trial Registry (http://www.chictr.org.cn, Registration number: ChiCTR-OCH-11001198).

## Introduction

As a standard treatment procedure for patients suffering from ACS and those undergoing PCI with stenting, dual antiplatelet therapy(DAPT) with clopidogrel in addition to aspirin, significantly reduces the risk of adverse cardiac events in patients.[1] However, the pharmacodynamic response to DAPT varies substantially among patients.[2]

Studies have reported the associations of several gene polymorphisms (*CYP2C19*2, CYP2C19*3, CYP2C9*2, PON1* Q192R and *ABCB1* C3435T) with cardiovascular outcomes in patients with ACS and those undergoing PCI.[3-8] However, there are meta-analysis studies failed to support the associations of CYP2C19 and cardiovascular events.[9, 10] In summary, these studies about the effects of single nucleotide variants (SNPs) on clinical outcomes were inconsistent and inconclusive. Meanwhile, most of the previous studies were conducted in western populations, Asians have very different genotype distributions of CYP2C19 and PON1 Q192R, in whom relevant studies based on large samples are scarce. Therefore, it’s very necessary to identify the association of genetic polymorphisms with cardiovascular events and offer valuable information for intervention in Han Chinese population. Moreover, previous studies on cardiac adverse events after PCI mainly focused on a few specific genotypes or genes.[3-10] Therefore, the majority of the hereditability in affecting cardiovascular events remains unexplained, and other important genetic determinants have yet to be identified. Genome-wide studies, such as whole exome sequencing, should be conducted to enhance the understanding of this research field.

In the present study, we combined whole exome and targeted sequencing to investigate the genetic factors underlying the major adverse cardiovascular events (MACE) among patients receiving clopidogrel and aspirin treatments after PCI. The logistic regression and Cox proportional hazard models were used to analyze 1961 samples in two independent cohorts with detailed clinical information. This process aimed to evaluate the previously reported cardiovascular outcome-related loci and discovered novel genes/alleles associated with the effect of treatment with clopidogrel and aspirin on MACE in the Han Chinese ethnic group. To date, machine learning methods have not been substantially applied to develop effectively predictive prognostic classifiers for adverse cardiovascular events. Thus, we developed SVM classifier to predict the occurrence possibility of MACE, as well as to provide evidence for therapeutic intervention of patients with ACS.

## Methods

### Study population and study design

In this study, all patients with acute coronary syndromes (ACS) undergoing PCI were obtained through Guangdong General Hospital in China from 2009 to 2012. These patients were treated with 12 months of dual antiplatelet therapy with clopidogrel in addition to aspirin following stent implantation. Patient information was collected based on inpatient and outpatient hospital visits, and telephone contacts with the patients or their family at 1, 6, 12 and 18 months following discharge. During the 18-months follow-up period, trained staff systematically recorded medical conditions of each patient to determine the occurrence of major adverse cardiovascular events (MACE). A physician adjudicated the end point through review of source documents obtained from medical records. The major adverse cardiovascular events (MACE) as a composite endpoint included cardiovascular death, myocardial infarction (MI), stroke (CT or MR scan confirmed) and repeated revascularization (RR). Repeat revascularization included target vessel revascularization-PCI, non–target vessel revascularization-PCI, and coronary artery bypass grafting (CABG).

We included 1961 patients who insisted on DAPT therapy for 12 months and had detailed baseline and follow-up information during 18-months follow-up periods. These samples were divided into two cohorts. In the discovery cohort, we selected 168 patients, of which 51 had MACE end point during 1-year follow-up period and 117 didn’t had any clinical events during 18-months follow-up periods. In the replication cohort, a total of 1793 patients were included and 1703 samples went into subsequent analysis after sample quality control, of which 123 had MACEs end point during 1-year follow-up period and 1580 didn’t had any clinical events during 18-months follow-up periods. This study has been registered in the Chinese Clinical Trial Registry (http://www.chictr.org.cn, Registration number: ChiCTR-OCH-11001198).

This study was approved by both the Guangdong general hospital ethics committee and the BGI ethics committee. All protocols were conducted in compliance with the Declaration of Helsinki, and explicit informed consent was obtained from all participants.

### Sample sequencing in two stages

We sequenced the whole exome of 168 patients in the discovery cohort. Genomic DNA for each sample was used to produce each exome captured library with the NimbleGen SeqCap EZ Exome (44M, Roche) array. Then each captured library was independently sequenced on the illumina Hiseq 2000 platform. Each sample was designed to get high quality bases with coverage of > 90×.

We sequenced the 6M targeted region of 1793 patients in replication cohort. The targeted region consisted of three parts. The first part was the top associated SNVs with P<0.05 in the discovery stage. The second part was top genes with P<0.05 in gene-based test in the discovery study. Gene-based analysis was performed using the Fast Association Tests (FAST) tool which includes a series of gene-based methods. Four algorithms, including (i) Gene-Wide Significance test (GWiS); (ii) MinSNP-p and MinSNP-gene; (iii) Versatile Gene-Based Test for Genome-wide Association (VEGAS); and (v) the Gates test (GATES), were used to test the gene-based association with MACE or bleeding events. The significant gene (p<0.05) in at least three of the four tests was selected for the targeted region design. The third part was 49 reported genes within pharmacokinetic and pharmadynamics pathway of clopidogrel, aspirin, statin or beta-blockers. All the three parts were combined and merged into 6M target regions. Then the targeted region for each sample was sequenced on Complete Genomics (CG) platform with high quality and coverage.

### Sample quality control

We required the samples to meet these criteria: (i) average sequencing depth ≥ 90× in exome sequencing stage; (ii) average sequencing depth ≥ 30× in targeted sequencing stage; (iii) genotype calling rate ≥ 90%; (iv) not existing population stratification by performing principal components analysis (PCA) analysis which were performed via the multidimensional scaling (MDS) procedure implemented in PLINK v 1.07 and (v) not be duplicates or first-degree relatives while evaluating pairwise by identity by descent (IBD). After quality control filtering, 90 samples from the targeted sequencing stage were excluded from subsequent analysis.

### Alignment, Variant calling and quality control

For illumina exome sequencing data, reads were mapped to human genome reference assembly (hg19, GRCh37) with SOAP2 and variants were detected by SOAPsnp. The high-quality illumina SNVs that we defined for each individual had to meet the following condition: sequenced quality ≥ 20, sequencing depth ≥ 8×, sequencing depth ≤ 500×, and depth of non-reference allele ≥ 4×.

For CG targeted sequencing data, sequence reads were also aligned to human reference genome hg19 and variations were detected using the CG analysis toolkit (CGATools) which is available at CG website (http://cgatools.sourceforge.net/).

After all initial SNV calls from illumina and CG platforms were generated, further filtering was performed to identify high-confidence SNVs. We required SNVs from discovery study and replication study to meet two conditions: (i) genotyping call rate ≥ 90%, (ii) MAF ≥ 0.01 and (iii) hardy–weinberg equilibrium (HWE) P > 1.0× 10^-6^.

To evaluate the data quality, we compared the genotypes from the sequencing data with the genotypes called from the genotyping arrays. In the discovery study, the average genotype concordance is 98.6% by comparing 5 genotypes overlapping in the exome sequencing data and genotyping array data in 126 samples (**Table S1**). In the replication study, the whole genome sequencing result of the YanHuang (YH) sample was obtained using CG. We evaluated the CG data quality by comparing the whole genome sequencing result and genotyping results in the YH sample and the genotype concordance was 99.5%.

### Statistical analysis

Analysis was performed using PLINK (version 1.07)[11] and R (version 3.2.3, http://www.R-project.org/). The demographic and clinical characteristics were summarized using counts (percentages) for the categorical variables (e.g., sex, previous MI, diabetes mellitus, etc.) and mean (standard deviation, SD) for the continuous variables (e.g., age, BMI, etc.). A univariate Cox proportional hazard model was used to calculate the significance by comparing the baseline demographic and clinical characteristics between the groups with and without MACE. In the discovery cohort, we applied the logistic regression to calculate the P values and odds ratio (OR) of SNPs on the clinical end points by adjusting for 17 variables, including the first four principal components PC1-PC4, 3 demographic (sex, age, BMI) and 10 clinical variables (e.g., Diabetes mellitus, hypertension, previous MI, etc.). In the replication cohort, we use the multivariate Cox proportional hazards regression to model the survive time and the incidence of MACE by adding SNPs and the same 17 adjustment variables as covariates to evaluate the hazard ratio (HR) and P value for each SNP. Finally, we also perform the multivariate Cox proportional hazards regression analysis to evaluate the associations of the genotypes with MACE in the combined data.

The Cox model is expressed by the hazard function denoted by h(t). Briefly, the hazard function can be interpreted as the risk of MACE at time t. It can be estimated as follow:

h(t)=h0(t)×exp(b1×1+b2×2+…+bpxp)

where,

- t represents the survival time of MACE, ranging from 0 to 12 months.
- h(t) is the hazard function determined by a set of p covariates (x1, x2,…, xp), p covariates were SNPs and the 17 adjustment variables in this study.
- the coefficients (b1, b2,…, bp) measure the impact (i.e., the effect size) of covariates.
- the term h0 is called the baseline hazard. It corresponds to the value of the hazard if all the xi are equal to zero (the quantity exp(0) equals 1). The ‘t’ in h(t) reminds us that the hazard may vary over time.

We fit the multivariate Cox proportional hazards regression model with coxph() function in survival package in R. The coxph function was written as:

>coxph(Surv(time, status) ∼ age + sex +… + genotypes, data = MACE)

Baseline variables and genetic variants explanation for MACE were calculated by using a regression-based approach as implemented in the SOLAR-Eclipse version 8.1.1 software (http://solar-eclipse-genetics.org/index.html).

Power analysis was used to investigate if we have enough power to detect the association SNPs in the sample size. R package survSNP (version 0.23.2) was used to evaluate the power to achieve a P value of 10^-6^ for association SNPs with different HR or MAF.

### Gene expression analysis

We mined four publically available genome-wide expression data sets from Gene Expression Omnibus (GEO) database in NCBI, including GSE27962[12], GSE48060[13], GSE7487[14] and GSE47495[15]. These datasets recorded gene expression data of cardiac remodeling after myocardial infarction or Myocardial Infarction-Induced Heart Failure. We compared the gene expression difference between case groups (mainly myocardial infarction (MI) operation) and control groups (sham operation) and used unpaired t-test to compute the significance. Genes with two-tailed P < 0.05 were considered to be differentially expressed between two groups.

### *ECHS1* plasma experiments

We randomly selected ACS patients with heart failure (HF) symptoms at NYHA stage II or less (n=61), stage III or IV (n=89) from an independent study cohort. *ECHS1* protein levels was measured in plasma sample using sandwich enzyme-linked immunosorbent assays (ELISA) (ECHS1 ELISA kit, Action-award Biotech co. Ltd., Guangzhou, China) and a Multiskan GO Microplate Reader (Thermo Scientific Inc., USA). The comparison of parameters and *ECHS1* protein levels between patients with HF at stage II or less(n=61) and patients at stage III or IV(n=89) was performed using the logistic regression analysis. Because the distribution of *ECHS1* protein levels was skewed, logarithmic transformation was performed before analysis.

### Predictive diagnostic power of clinical factors and genetic variants

This study used the support vector machine (SVM) method[16] to predict whether MACE occurred in a patient during the 18-month follow-up. we selected six features: 1) all clinical factors; 2) all genetic variants; 3) combined all clinical factors and all genetic variants; 4) 3 significant clinical factors, including age, hypertension and creatinine; 5) 27 significant genetic variants with P<10-4; 6) combined 3 significant clinical factors and 27 significant genetic variants, and constructed the classifier for each feature to estimate the diagnostic effectiveness of each feature. For each feature, the e1071 package in R (www.r-project.org) was used to construct the classifier via the SVM function with the nonlinear radial basis function as the kernel. During the training process, we use five-fold cross-validation technique to verify the predictive power of each classifier. All 1961 samples are randomly divided into almost the same number of 5 subsets, four subsets are used as training sets and the remaining one subset is used as a test set each time, the process is repeated five times so that every subset is used as a test set once. The performance of the classifier is the average of the 5 sets performances. Finally, five indices were utilized: sensitivity, specificity, positive predictive value, negative predictive value, and area under the receiver operating characteristic (ROC) curve. For each feature, the sensitivity and specificity of its classifier as a predictor of MACE was evaluated by the ROC curves using the area under the curve (AUC) as a measure of diagnostic effectiveness. Finally, the feature of combined 3 significant clinical factors and 27 significant genetic variants were kept to develop the classifier with the best predictive diagnostic effectiveness of MACE.

## Results

### Characteristics of the Patients

Two independent cohorts were recruited and comprised 1961 patients with ACS who underwent PCI and treated with clopidogrel and aspirin for 6 to 12 months in accordance with the consensus guidelines. After performing sample quality control, 168 patients were included in the discovery study and 1703 patients were remained in the replication study (**Table S2 and S3**). For the total 1871 patients, the average age was 63.3 (±10.7) years, 357 (19.1%) were women, 174 (9.3%) had a MACE endpoint. **Table 1** shows the clinical characteristics of 174 cases with MACE and 1697 controls without MACE. MACE was associated with increasing age (65.84 ± 10.17 vs. 63.07 ± 10.74, P = 0.001), hypertension (68% vs. 56%, P = 0.001), and high creatinine (100.9 ± 85.96 vs. 87.03 ± 44.84, P = 0.001) (**Table 1**). These combined clinical variables explained 5.80% of occurrence of the MACE, thus suggesting potentially substantial genetic contribution.

**Table 1.** Baseline characteristics of case-control patients.

| Characteristics | With<br>MACE<br>(n=174) | Without<br>MACE<br>(n=1697) | P value | Variants<br>explanation |
| --- | --- | --- | --- | --- |
| Age – yrs, mean (±s.d.) | 65.84(±10.17) | 63.07(±10.74) | 0.001** | 0.94% |
| Sex, Men – no. (%) | 139(80) | 1375(81) | 0.71 |  |
| BMI –kg/m<sup>2</sup>, mean (±s.d.) | 24.14(±3.08) | 23.89(±3.18) | 0.58 | |
| **Risk factors, n (%)** |  |  |  |  |
| Previous MI | 67(39) | 627(37) | 0.65 |  |
| Diabetes mellitus | 59(34) | 468(28) | 0.08 |  |
| Hypertension | 119(68) | 944(56) | 0.001** | 0.87% |
| **Medications used before event, n (%)** |  |  |  |  |
| ACEI_ARB | 150(86) | 1385(82) | 0.17 |  |
| BBi | 155(89) | 1475(87) | 0.5 |  |
| CCB | 44(25) | 364(21) | 0.23 |  |
| PPI | 91(52) | 786(46) | 0.12 |  |
| Statins | 170(98) | 1648(97) | 0.89 |  |
| **Clinical laboratory characteristics, mean (s.d.)** |  |  |  |  |
| HDLC, mmol/L | 0.98(0.24) | 0.98(0.26) | 0.66 |  |
| LDLC, mmol/L | 2.59(0.92) | 2.7(0.99) | 0.13 |  |
| Triglycerides, mmol/L | 1.51(0.94) | 1.57(1.21) | 0.5 |  |
| HbA1c, % total haemoglobin | 6.39(0.9) | 6.59(1.42) | 0.12 |  |
| ALT, U/L | 38.09(56.35) | 32.2(32.85) | 0.06 |  |
| AST, U/L | 33.06(33.73) | 39(57.85) | 0.21 |  |
| CREA, μmol/L | 100.9(85.96) | 87.03(44.84) | 0.001** | 0.73% |
| CK, U/L | 106.68(80.86) | 115.32(86.89) | 0.28 |  |
| CKMB, U/L | 6.50(4.84) | 6.75(4.68) | 0.68 |  |

P values were calculated by univariate Cox regression analysis. P < 0.05 was considered a statistically significant difference.
BMI denotes Body-mass index; MI myocardial infarction; ACEI angiotensin-converting-enzyme inhibitor; ARB angiotensin receptor blocker; BBi $\beta$ -blockers inhibitors; CCB calcium channel blockers; PPI proton pump inhibitors; HDLC highdensity lipoprotein cholesterol; LDLC low-density lipoprotein cholesterol; HbA1c hemoglobin A1c; ALT alanine aminotransferase; AST aspartate aminotransferase; CREA creatinine; CK creatine kinase and CKMB creatine kinase MB.
Signif. codes: 0 '\*\*\*\*' 0.001 '\*\*\*' 0.01 '\*\*' 0.05 '.' 0.1 ' ' 1

### Two-stage association study

The study design and total workflow is shown in **Figure 1**. In the discovery study, we deep-sequenced the whole exome of the 168 ACS patients with a mean coverage of approximately 210× (**Table S4 and Figure S1**). All cases with MACE and controls without MACE were ethnically and genetically well matched (**Figure S2**). A total of 127,834 SNPs passed the quality control for the single-variant association analysis. Logistic regression analysis determined 6268 SNPs associated with MACE with P < 0.05 adjusting for the covariates **(Figures S3**). Gene-based association analysis identified 408 genes associated with MACE at the P < 0.05 level (**Table S5**).

**Figure 1.**
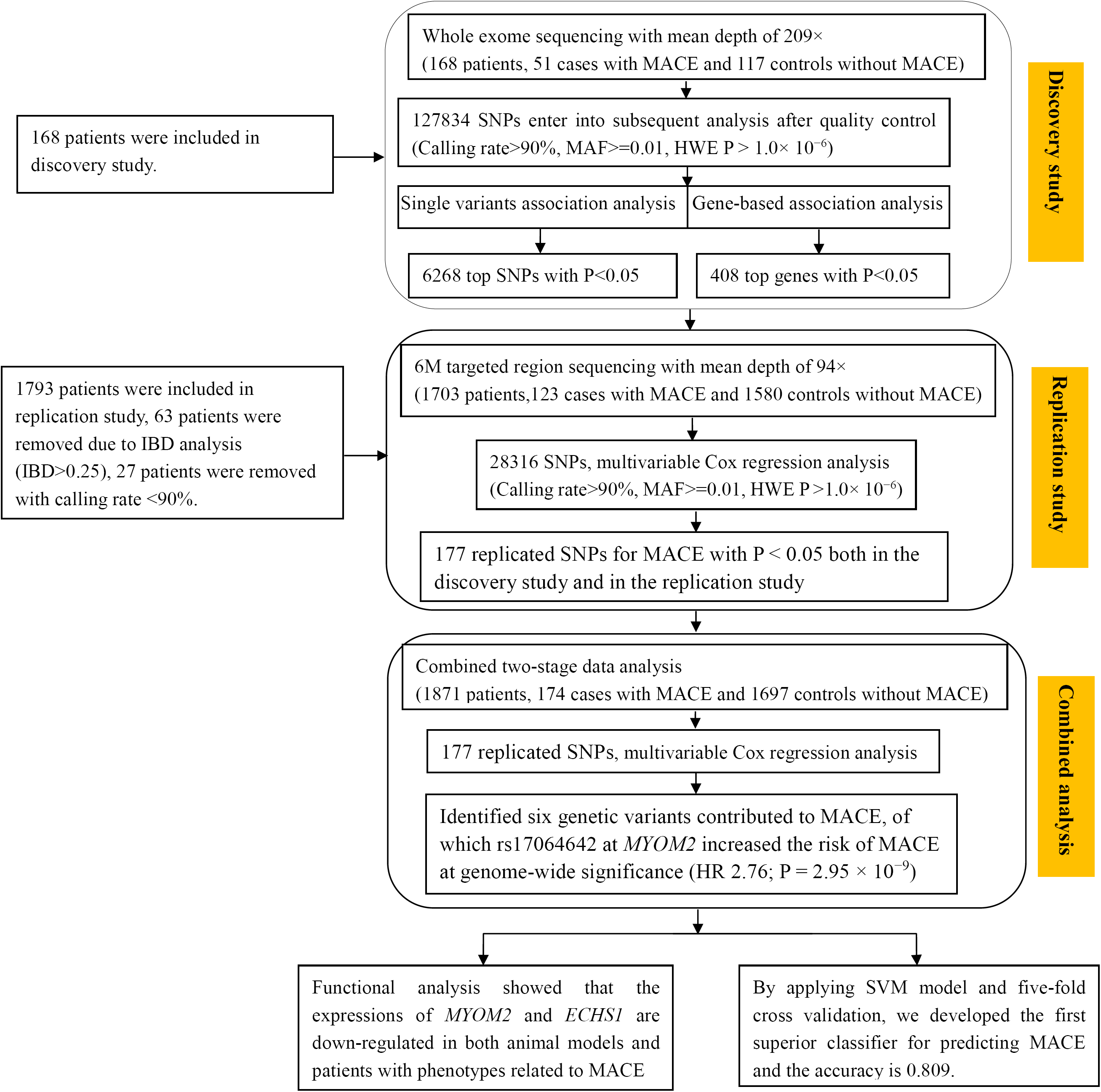
Description of the study design. First, we performed exome sequencing of 168 patients and 51 had MACE end point. After quality filtering, a total of 127834 variants was subjected to single variant association analysis and 6268 variants showed nominal association (P < 0.05). Gene-based association analyses identified 408 genes associated with MACE (P < 0.05). As validation, the 6M targeted region including 6268 top SNPs and 408 top genes were further analyzed in additional 1793 patients through multivariable Cox regression analysis. A total of 177 replicated SNPs in validation datasets went into combined data analysis. Finally, we identified six genetic variants contributed to MACE after multiple test correlation (P < 1.0 × 10^-6^). Then, we performed functional analysis on the six significant SNPs or genes, further we developed the first superior classifier for predicting MACE.

To further replicate the identified associations in the discovery cohort and increase the statistical power of the present study, we performed a replication analysis in an independent cohort of 1703 patients using targeted sequencing. In the replication analysis, a total of 6M targeted regions consisting of SNP/gene associations discovered above and previously reported, were sequenced with approximately 94× coverage for each individual (**Table S5 and Figure S1**). After performing quality control on the samples and variants, 28,316 SNPs entered the subsequent analyses (**Figures S3)**. Finally, we found 177 replicated SNPs for MACE with P < 0.05 both in the discovery and replication studies. For the 177 replicated signals, we performed multivariate cox regression analyses in the combined data set from 1871 samples (i.e., 174 MACE cases and 1697 controls) and adjusted for multiple covariates.

### Identify genetic variants associated with MACE

We identified one genotype rs17064642 at *MYOM2* was associated with MACE and achieved genome-wide significance (P = 2.95 × 10^-9^) (**Table 2**). Carriers of the CC/CT genotype had higher MACE occurrence rates during the 18 months of follow up compared with noncarriers (17.8% vs. 8.1%; HR, 2.76; 95% CI, 1.98–3.87) (**Figure 2**). Rs17064642 has moderate LD of r^2^ = 0.57 (East Asians) with the missense SNP rs34823600 and these two SNPs are located only 16bp apart. Rs17064642 also coincides with enhancer markers (H3K4me1_Enh and H3K27ac_Enh) in four tissues, especially in heart cell types (right Atrium, left Ventricle and right Ventricle), suggesting this locus may function as an enhancer in heart tissue (**Figure S5**).

**Table 2.**
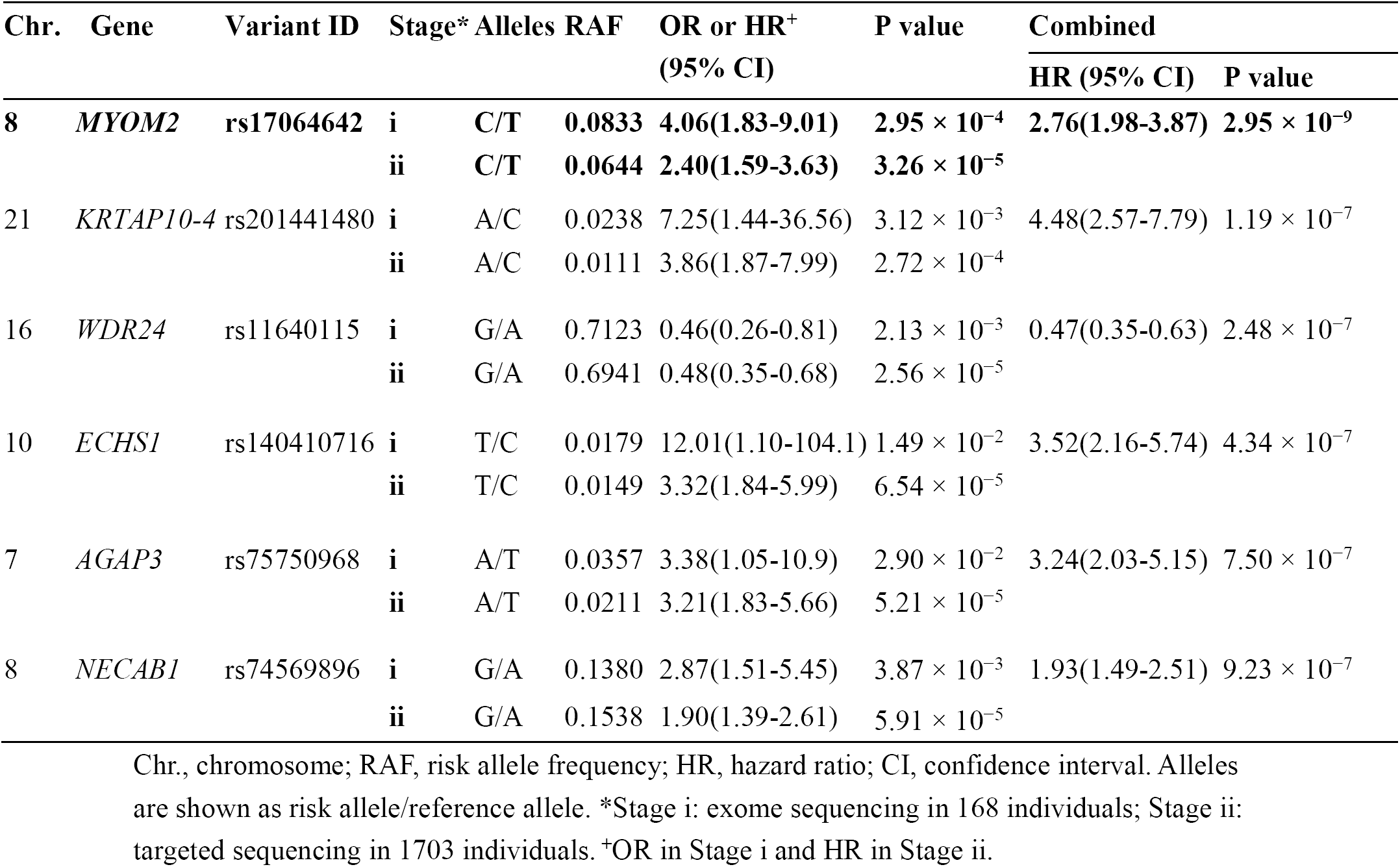
Identifying six genetic variants contributed to MACE.

**Figure 2.**
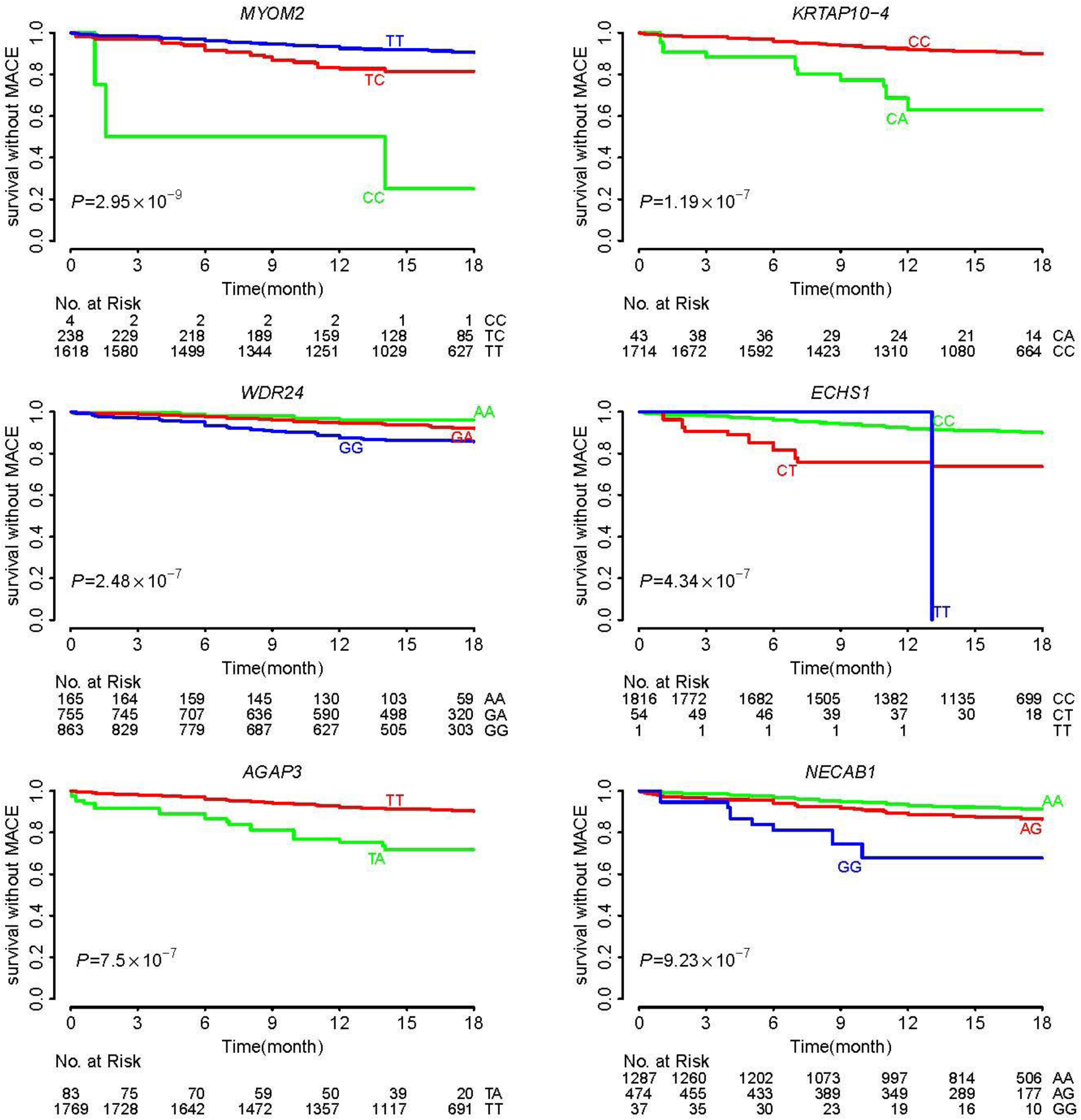
Event-Free survival over 18 months of follow-up in 1871 patients with ACS. Cumulative probabilities of survival without MACE according to gene polymorphisms: *MYOM2* (rs17064642), *KRTAP10-4* (rs201441480), *WDR24* (rs11640115), *ECHS1* (rs140410716), *AGAP3* (rs75750968) and *NECAB1* (rs74569896).

We also observed five additional significant associations after correction for multiple testing (P=0.05/28316=1.0 × 10^-6^), including *KRTAP10-4* (rs201441480), *WDR24* (rs11640115), *ECHS1* (rs140410716), *AGAP3* (rs75750968) and *NECAB1* (rs74569896) (**Table 2**).Rs201441480 at *KRTAP10-4* as one missense alteration was predicted to be damaging by Polyphen2 and significantly increased the risk of MACE (HR 4.48, P = 1.19 × 10^-7^). At *WDR24*, we identified two SNPs associated with MACE, rs11640115 (HR 0.47, P=2.48 × 10^-7^) and rs763053 (HR 0.52, P=3.02 × 10^-6^), which were in complete LD (D′ = 1, r^2^ = 1). According to public GTEx [17] databases, both rs11640115 and rs763053 at *WDR24* showed significant eQTL associations with gene *WDR90* (P=1.16 × 10^-6^) in the heart tissues, including heart left ventricle and heart atrial appendage. Evidence from GSE48060[13] dataset confirmed that *WDR90* transcription was associated with long-term recurrent events following first-time MI. Similarly, we also identified two SNPs at *NECAB1* associated with MACE, rs74569896 (HR 1.93, P=9.23 × 10^-7^) and rs73694346 (HR 1.72, P=4.86 × 10^-5^), which were in strong LD (D′ = 0.96, r^2^= 0.92). SNP rs75750968 locates in gene *AGAP3*, which relates to GTP binding and GTPase activator activity.

Together, we discovered six novel genetic variants associated with MACE and achieved an average of 87% power for the six SNPs, of which, 97.5% power for rs17064642 at *MYOM2* (**Table S6 and Figure S4**). **Figure 2** showed cumulative proportions of individuals without MACE over 18 months of follow-up under the six polymorphisms.

### Biological implications of variants associated with MACE

To further explore the functional evidence of the six genetic predispositions that contribute to MACE, we searched all literatures referring to the six genes and cardiovascular diseases in PubMed database. Meanwhile, we also mined several publicly available genome-wide expression data sets from the GEO database,[12-15] which recorded cardiac remodeling data after myocardial infarction (MI). We found two of six genes, *MYOM2* and *ECHS1*, showed abundant evidences of decreased expression in cases with adverse cardiac events, either in previous reports or in our data analysis. M-Protein (myomesin-2) encoded by *MYOM2* or total myomesin is down-regulated either in cardiac hypertrophy in rats,[18] in acute myocardial infarction (AMI) patients,[19] or in chronic heart failure.[20] Similarly, *ECHS1,* as an ischaemic post-conditioning (PostC) modified protein, showed significantly decreased expression in the cardiac remodeling group after MI compared with the sham group, by analyzing the two different groups’ expression data from GEO database GSE7487[14] (12,930 ± 337 vs. 16,360 ± 466, P = 5.46 × 10^-6^) and GSE47495[15] (11.89 ± 0.11 vs. 12.13 ± 0.04, P = 7.28 × 10^-4^, **Table S7**). Meanwhile, we compared the plasma *ECHS1* protein levels in ACS patients with HF symptoms at New York Heart Association (NYHA) stage III or IV (n=89) to those with HF symptoms at NYHA stage II or less (n=61). We found advanced HF patients showed significantly decreased levels of *ECHS1* protein (215.18±115.67 for stage II or less vs. 161.84±76.67 for stage III or stage IV, P = 0.0012) (**Table S8 and Figure S6**).

### Predictive accuracy of clinical factors and genetic variants for MACE

To develop a prognostic classifier for MACE, we investigated their predictive power to MACE based on different combinations of clinical factors and genetic variants. The predictive accuracy of single variable (i.e., clinical factor or genetic variant) for MACE, as determined by the receiver operating characteristic (ROC) curves, was insufficient with AUC varying between 0.4 and 0.6 (**Figure S7**). The predictive effectiveness of all clinical factors, all genetic variants, and combined all clinical factors and all genetic variants were 0.673, 0.683 and 0.684, respectively (**Figure 3**). However, the predictive power of the combined significant clinical factors and significant genetic variants for MACE was substantially increased compared with the single variable or all variables. Meanwhile, when we used the combined feature of 3 significant clinical factors (age, hypertension, creatinine) and 27 significant genetic variants showing suggestive significance with P < 10^-4^ (**Table S9**), the AUC for the MACE classifier was 0.809 and achieved best performance (**Figure 3**). In addition, the predictive power did not increase while adding more clinical factors or genetic variants to the analysis. Thus, the present study developed the first superior classifier for predicting MACE by selecting the most informative factors that independently contributed to MACE.

**Figure 3.**
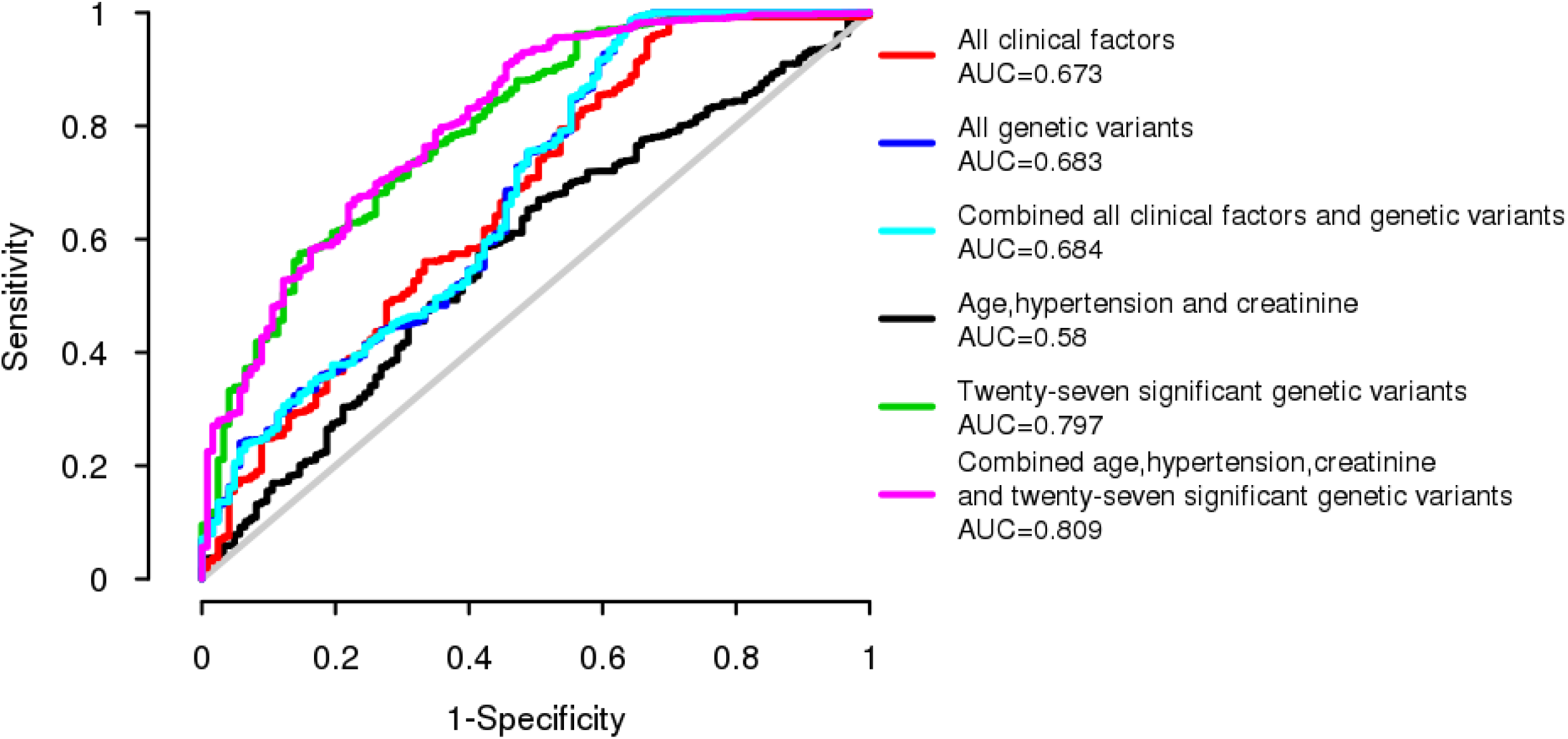
Receiver operating characteristic (ROC) curves for predictive features of MACE. ROC curves for six features: 1) all clinical factors; 2) all genetic variants; 3) combined all clinical factors and all genetic variants; 4) 3 significant clinical factors, including age, hypertension and creatinine; 5) 27 significant genetic variants with P<10-4; 6) combined 3 significant clinical factors and 27 significant genetic variants, and we constructed the classifier for each feature to estimate the diagnostic effectiveness of each feature for MACE.

## Discussion

To our knowledge, the present study is the first genome-wide and large-scale association analysis that integrates whole exome and targeted sequencing to identify novel genetic variants associated with MACE in patients with clopidogrel and aspirin treatment after PCI. By combining the two-stage sequencing data, we investigated five commonly reported SNPs[4, 6-8, 21-23] and confirmed *PON1* (HR 1.45; P = 0.003) but not *CYP2C19* genetic variants contributed to cardiovascular outcomes in Han Chinese patients (**Table S10**). Moreover, we identified six novel genetic variants associated with MACE: *MYOM2* (rs17064642), *KRTAP10-4* (rs201441480), *WDR24* (rs11640115), *ECHS1* (rs140410716), *AGAP3* (rs75750968) and *NECAB1* (rs74569896).

Among the six genes contributed to MACE, we find the expression levels of *MYOM2* and *ECHS1* are evidently down-regulated in cases subjected to adverse cardiac events compared with normal controls. Rs17064642 at *MYOM2* increased the risk of MACE and achieved genome-wide significance (HR, 2.76; 95% CI, 1.98–3.87; P = 2.95 × 10^-9^). *MYOM2* encodes the M-protein, which is also known as Myomesin-2. Myomesin-2 is the primary myosin M-band cross-linking protein and binds titin in a complex with obscurin/obs1. The protein is key to normal heart function, as evidenced by associations between heart failure and low expression. Myomesin-2 showed decreased expression in multiple heart diseases or heart attack. An animal model of cardiac hypertrophy driven by the thyroid hormone (T3) in rats showed that the *MYOM2* gene is severely down-regulated by T3 and M-protein deficiency causes significant contractile dysfunction (P < 0.05), thereby demonstrating the importance of M-protein (*MYOM2*) for normal cardial function.[18] In addition, it has been reported that myomesin (encoded by both *MYOM1* and *MYOM*2) protein levels decrease in acute ischemia and in chronic heart failure.[20] A proteomic analysis agrees with previous reports that level of myomesin-2 in cardiac tissue is decreased in AMI patients (n=10) compared with control cases (n=11).[19] Therefore, we speculated that variants in *MYOM2* affected the expression levels of M-protein and the decreased expression of M-protein causes significant contractile dysfunction involving in cardiac function, thereby causing the occurrence of MACE.

The *ECHS1* protein (short-chain enoyl-CoA hydratase) is a multifunctional mitochondrial enzyme with several functions in β-oxidation of short- and medium-chain fatty acids, as well as in isoleucine and valine metabolism. A previous study reported that the mitochondrial protein *ECHS1* could regulate cellular ATP consumption/production and influence the defense response to myocardial ischaemic stress.[24] Haack et al. reported that *ECHS1* deficiency causes mitochondrial encephalopathy with cardiac involvement.[25] The *ECHS1* deficiency can induce β-oxidation defect and impaire valine oxidation. The aforementioned authors speculated that the β-oxidation defect and block in L-valine metabolism, with accumulation of toxic methacrylyl-CoA and acryloyl-CoA, contribute to the disorder. Moreover, *ECHS1* showed significantly decreased expression in the cardiac remodeling group after MI compared with the sham group in two GEO datasets GSE7487 and GSE47495. Since cardiac remodeling has a clinical consequence of the heart failure (HF), blood *ECHS1* protein concentration may correlate with the presentation of HF. As expected, we confirmed *ECHS1* contributed to an ischemic heart failure in ACS patients by gene expression experiment. Thus, we assumed that the SNP rs140410716 G489A increases MACE risk (HR 3.52; 95% CI 2.16–5.74, P = 4.34 × 10^-7^) by down-regulating the expression of *ECHS1* in MI, which could reduce the cellular ATP consumption/production and cause MACE or cardiac failure as a result of mitochondrial dysfunction. Combining all these evidences, we speculate that *ECHS1* deficiency causes contractile dysfunction of the cardiac muscle and dysfunction of the mitochondrial energy metabolism, and eventually leads to MACE.

Apart from identifying the novel genotypes and genes associated with MACE, we improved the prediction of MACE by developing a novel classifier for MACE. To date, no comprehensive and complete genetic makers for an improved prognostic classification of MACE is available. The developed classifier can combine the clinical characteristics and multiple independently informative genotypes to predict MACE and achieved high accuracy (AUC=0.809).

In conclusion, we provide here the first genome-wide, large-scale association analysis on ACS patients receiving clopidogrel and aspirin treatment after PCI. We successfully identify six new and independent genetic predispositions for MACE and discover 12 variants associated with cardiovascular death, MI, stroke or RR. Of them, two variants reached genome-wide significance: Rs17064642 at *MYOM2* increased the risk of MACE and rs117279588 at *ATP6V1A* increased the risk of stroke. We find these variants may regulate the function of nearby genes, and the expressions of *MYOM2* and *ECHS1* are down-regulated in both animal models and patients with phenotypes related to MACE. These findings will provide clinicians with potential biomarkers for an improved prediction of MACE or cardiovascular death, MI, stroke or RR, and provide new insight on the therapeutics of ACS.

## Author contributions

S.L.Z. and X.L. conceived of and designed the research. X.M.L., H.S.X., and H.Q.X. managed the project. X.M.L. drafted the initial manuscript. S.L.Z., Q.S.G., J.Y.C. and X.Y.L. conducted sample selection and data management, R.G., P.P.J., H.M.Y., J.W., X.Q.Z. and X.X. generated the sequence data. X.M.L., H.S.X., H.Q.X., W.H.M., Z.S., F.Y. and T.Z. analyzed and interpreted the data. X.M.L., H.S.X., H.Q.X. and W.H.M. did statistical analyses. X.M.L., H.S.X., H.Q.X., Q.S.G., X.L. and S.L.Z. made critical revisions to the manuscript for important intellectual content. Each author is responsible for the content and the writing of the paper. The authors declare that they have no competing interests.

## Acknowledgments

We gratefully acknowledge Guangdong general hospital for sample collection and thank colleagues at BGI-Shenzhen for DNA extraction, library construction, sequencing, and discussions. The research was supported by the National key R&D program (No. 2017YFC0909301, 2016YFC0905003), National Nature Science Foundation of China (No. 81673514, 81373486), Science and Technology Development Projects of Guangdong Province, China (No. 2013B021800157, 2016B090918114), and Science and Technology Development Projects of Guangzhou, Guangdong, China (No. 201510010236).

## References

1. Zeymer U, Gitt A, Junger C, Bauer T, Heer T, Koeth O, Mark B, Zahn R, Senges J, Gottwik M: Clopidogrel in addition to aspirin reduces in-hospital major cardiac and cerebrovascular events in unselected patients with acute ST segment elevation myocardial. Thromb Haemost 2008, 99(1):155–160.

2. Gurbel PA, Bliden KP, Hiatt BL, O’Connor CM: Clopidogrel for coronary stenting: response variability, drug resistance, and the effect of pretreatment platelet reactivity. Circulation 2003, 107(23):2908–2913.

3. Mega JL, Simon T, Collet JP, Anderson JL, Antman EM, Bliden K, Cannon CP, Danchin N, Giusti B, Gurbel P et al: Reduced-function CYP2C19 genotype and risk of adverse clinical outcomes among patients treated with clopidogrel predominantly for PCI: a meta-analysis. JAMA 2010, 304(16):1821–1830.

4. Shuldiner AR, O’Connell JR, Bliden KP, Gandhi A, Ryan K, Horenstein RB, Damcott CM, Pakyz R, Tantry US, Gibson Q et al: Association of cytochrome P450 2C19 genotype with the antiplatelet effect and clinical efficacy of clopidogrel therapy. JAMA 2009, 302(8):849–857.

5. Mega JL, Close SL, Wiviott SD, Shen L, Hockett RD, Brandt JT, Walker JR, Antman EM, Macias W, Braunwald E et al: Cytochrome P-450 Polymorphisms and Response to Clopidogrel. New England Journal of Medicine 2009, 360(4):354–362.

6. Bouman HJ, Schomig E, van Werkum JW, Velder J, Hackeng CM, Hirschhauser C, Waldmann C, Schmalz HG, ten Berg JM, Taubert D: Paraoxonase-1 is a major determinant of clopidogrel efficacy. Nat Med 2011, 17(1):110–116.

7. Chen Y, Huang X, Tang Y, Xie Y, Zhang Y: Both PON1 Q192R and CYP2C19*2 influence platelet response to clopidogrel and ischemic events in Chinese patients undergoing percutaneous coronary intervention. Int J Clin Exp Med 2015, 8(6):9266–9274.

8. Mega JL, Close SL, Wiviott SD, Shen L, Walker JR, Simon T, Antman EM, Braunwald E, Sabatine MS: Genetic variants in ABCB1 and CYP2C19 and cardiovascular outcomes after treatment with clopidogrel and prasugrel in the TRITON-TIMI 38 trial: a pharmacogenetic analysis. Lancet 2010, 376(9749):1312–1319.

9. Holmes MV, Perel P, Shah T, Hingorani AD, Casas JP: CYP2C19 genotype, clopidogrel metabolism, platelet function, and cardiovascular events: a systematic review and meta-analysis. JAMA 2011, 306(24):2704–2714.

10. Bauer T, Bouman HJ, van Werkum JW, Ford NF, ten Berg JM, Taubert D: Impact of CYP2C19 variant genotypes on clinical efficacy of antiplatelet treatment with clopidogrel: systematic review and meta-analysis. BMJ 2011, 343:d4588.

11. Purcell S, Neale B, Todd-Brown K, Thomas L, Ferreira MA, Bender D, Maller J, Sklar P, de Bakker PI, Daly MJ et al: PLINK: a tool set for whole-genome association and population-based linkage analyses. Am J Hum Genet 2007, 81(3):559–575.

12. Kuster DW, Merkus D, Kremer A, van Ijcken WF, de Beer VJ, Verhoeven AJ, Duncker DJ: Left ventricular remodeling in swine after myocardial infarction: a transcriptional genomics approach. Basic Res Cardiol 2011, 106(6):1269–1281.

13. Suresh R, Li X, Chiriac A, Goel K, Terzic A, Perez-Terzic C, Nelson TJ: Transcriptome from circulating cells suggests dysregulated pathways associated with long-term recurrent events following first-time myocardial infarction. J Mol Cell Cardiol 2014, 74:13–21.

14. Lin RC, Weeks KL, Gao XM, Williams RB, Bernardo BC, Kiriazis H, Matthews VB, Woodcock EA, Bouwman RD, Mollica JP et al: PI3K(p110 alpha) protects against myocardial infarction-induced heart failure: identification of PI3K-regulated miRNA and mRNA. Arterioscler Thromb Vasc Biol 2010, 30(4):724–732.

15. Tulacz D, Mackiewicz U, Maczewski M, Maciejak A, Gora M, Burzynska B: Transcriptional profiling of left ventricle and peripheral blood mononuclear cells in a rat model of postinfarction heart failure. BMC Med Genomics 2013, 6:49.

16. Vapnik VN: An overview of statistical learning theory. IEEE Trans Neural Netw 1999, 10(5):988–999.

17. Consortium GT: Human genomics. The Genotype-Tissue Expression (GTEx) pilot analysis: multitissue gene regulation in humans. Science 2015, 348(6235):648–660.

18. Rozanski A, Takano AP, Kato PN, Soares AG, Lellis-Santos C, Campos JC, Ferreira JC, Barreto-Chaves ML, Moriscot AS: M-protein is down-regulated in cardiac hypertrophy driven by thyroid hormone in rats. Mol Endocrinol 2013, 27(12):2055–2065.

19. Kakimoto Y, Ito S, Abiru H, Kotani H, Ozeki M, Tamaki K, Tsuruyama T: Sorbin and SH3 Domain-Containing Protein 2 Is Released From Infarcted Heart in the Very Early Phase: Proteomic Analysis of Cardiac Tissues From Patients. Journal of the American Heart Association 2013, 2(6):e000565–e000565.

20. Hein S, Kostin S, Heling A, Maeno Y, Schaper J: The role of the cytoskeleton in heart failure. Cardiovasc Res 2000, 45(2):273–278.

21. Brandt JT, Close SL, Iturria SJ, Payne CD, Farid NA, Ernest CS, 2nd, Lachno DR, Salazar D, Winters KJ: Common polymorphisms of CYP2C19 and CYP2C9 affect the pharmacokinetic and pharmacodynamic response to clopidogrel but not prasugrel. J Thromb Haemost 2007, 5(12):2429–2436.

22. Simon T, Verstuyft C, Mary-Krause M, Quteineh L, Drouet E, Meneveau N, Steg PG, Ferrieres J, Danchin N, Becquemont L et al: Genetic determinants of response to clopidogrel and cardiovascular events. N Engl J Med 2009, 360(4):363–375.

23. Li XQ, Ma N, Li XG, Wang B, Sun SS, Gao F, Mo DP, Song LG, Sun X, Liu L et al: Association of PON1, P2Y12 and COX1 with Recurrent Ischemic Events in Patients with Extracranial or Intracranial Stenting. PLoS One 2016, 11(2):e0148891.

24. Zhu SG, Xi L, Kukreja RC: Type 2 diabetic obese db/db mice are refractory to myocardial ischaemic post-conditioning in vivo: potential role for Hsp20, F1-ATPase delta and Echs1. J Cell Mol Med 2012, 16(4):950–958.

25. Haack TB, Jackson CB, Murayama K, Kremer LS, Schaller A, Kotzaeridou U, de Vries MC, Schottmann G, Santra S, Buchner B et al: Deficiency of ECHS1 causes mitochondrial encephalopathy with cardiac involvement. Ann Clin Transl Neurol 2015, 2(5):492–509.

